# A chromosome-scale assembly for ‘d’Anjou’ pear

**DOI:** 10.1101/2023.08.22.554305

**Authors:** Alan Yocca, Mary Akinyuwa, Nick Bailey, Brannan Cliver, Harrison Estes, Abigail Guillemette, Omar Hasannin, Jennifer Hutchison, Wren Jenkins, Ishveen Kaur, Risheek Rahul Khanna, Madelene Loftin, Lauren Lopes, Erika Moore-Pollard, Oluwakemisola Olofintila, Gideon Oluwaseye Oyebode, Jinesh Patel, Parbati Thapa, Martin Waldinger, Jie Zhang, Qiong Zhang, Leslie Goertzen, Sarah B. Carey, Heidi Hargarten, James Mattheis, Huiting Zhang, Teresa Jones, LoriBeth Boston, Jane Grimwood, Stephen Ficklin, Loren Honaas, Alex Harkess

## Abstract

Cultivated pear consists of several *Pyrus* species with *P. communis* (European pear) representing a large fraction of worldwide production. As a relatively recently domesticated crop and perennial tree, pear can benefit from genome-assisted breeding. Additionally, comparative genomics within Rosaceae promises greater understanding of evolution within this economically important family. Here, we generate a fully-phased chromosome-scale genome assembly of *P. communis* cv. ‘d’Anjou’. Using PacBio HiFi and Dovetail Omni-C reads, the genome is resolved into the expected 17 chromosomes, with each haplotype totalling nearly 540 Megabases and a contig N50 of nearly 14 Mb. Both haplotypes are highly syntenic to each other, and to the *Malus domestica* ‘Honeycrisp’ apple genome. Nearly 45,000 genes were annotated in each haplotype, over 90% of which have direct RNA-seq expression evidence. We detect signatures of the known whole-genome duplication shared between apple and pear, and we estimate 57% of d’Anjou genes are retained in duplicate derived from this event. This genome highlights the value of generating phased diploid assemblies for recovering the full allelic complement in highly heterozygous crop species.

## Introduction

*Pyrus* L. is a genus in the family Rosaceae (subfamily Maloideae) comprising cultivated and wild pears. *Pyrus* is divided into two broad categories, the European and Asian pears, with their divergence estimated around 3-6 million years ago (Wu et al. 2018). At least 28 species of *Pyrus* and 10 naturally occurring interspecific crosses are now found in Western and Eastern Asia, Europe, North Africa, and the Middle East. In 2021, the pear’s value of utilized production in the United States reached $353 million (United States Department of Agriculture National Agricultural Statistics Service 2023). This makes pear one of the most cultivated pome fruits worldwide. One of the most important North American varieties of pear, the Anjou, also known as the Beurre d’Anjou or simply d’Anjou (*Pyrus communis* cv. ‘d’Anjou’), is thought to have originated in Belgium, named for the Anjou region of France.

Over the last decade, several pear genomes have been sequenced and assembled using a variety of technologies. The first *Pyrus* genome sequenced in 2012 was the most commercially important Asian pear *P. bretschneideri* Rehd. cv. ‘Dangshansuli’, using a combination of BAC-by-BAC sequencing and mate-pair Illumina sequencing (Wu et al. 2013). Following that, European pear (*P. communis* cv. ‘Bartlett’) was sequenced using Roche 454 (Chagné et al. 2014). In 2019, the *P. communis* genome was updated by sequencing the double-haploid ‘Bartlett’ cultivar using PacBio long reads and high-throughput chromosome conformation capture (Hi-C) technology (Linsmith et al. 2019). This assembly helped uncover duplicated gene models in previous assemblies that over-assembled heterozygous regions. However, being a double-haploid, it still lacked an entire parental complement. A draft assembly and annotation for *P. communis* cv. ‘d’Anjou’ was generated recently (H. Zhang et al. 2022), which was carefully annotated and revealed systematic differences in gene annotations across Rosaceae genomes. However, this assembly was also not phased, lacking information on allelic variants. Genomes are currently available for five of twenty-two *Pyrus* species in the Genome Database for Rosaceae (GDR; https://www.rosaceae.org/organism/26137), and for only a few of the thousands of recognized cultivars (J. Li et al. 2022).

Here, we sequenced and assembled a chromosome-scale reference genome for *Pyrus communis* cv. ‘d’Anjou’ using PacBio HiFi and Dovetail Omni-C sequencing. This genome was assembled as part of a semester-long undergraduate and graduate genomics course under the American Campus Tree Genomes (ACTG) initiative, where undergraduate and graduate students assemble, annotate, and publish culturally and economically valuable tree species. Here we present a haplotype-resolved, chromosome-scale assembly and annotation of ‘d’Anjou’ pear, place it in a phylogenetic context with other Rosaceae species, and show evidence of an ancient whole-genome duplication (WGD) event shared by cultivated apple and pear.

## Methods

### Genome sequencing and assembly

DNA was isolated from young leaf tissue using a standard CTAB approach (Doyle and Doyle 1987). Illumina TruSeq DNA PCR-free libraries were constructed from 1 μg of input DNA and sequenced on an Illumina NovaSeq6000 at HudsonAlpha Institute for Biotechnology. Raw reads were assessed for quality using FASTQC v0.11.9 (Andrews et al. 2010). Then, low quality reads were filtered out of the raw data by using *fastp* v0.12.4, allowing the generation of a statistical report with MultiQC 1.13.dev0 (Ewels et al. 2016). Nuclear genome size and ploidy was estimated using jellyfish v2.2.10 ((Ranallo-Benavidez, Jaron, and Schatz 2020; Marçais and Kingsford 2011)) to count k-mers, and visualized in GenomeScope2.0 (Ranallo-Benavidez, Jaron, and Schatz 2020; Marçais and Kingsford 2011). For PacBio HiFi sequencing, approximately 20 grams of young leaf tissue from a ‘d’Anjou’ pear clone was collected and flash-frozen in liquid nitrogen. High molecular weight DNA was isolated from the young leaf tissue using a Circulomics Nanobind Plant Nuclei Big DNA kit (Baltimore, MD), with 4 g of input tissue and a 2 hour lysis. DNA was tested for purity via spectrophotometry, quantified by Qubit dsDNA Broad Range, and size selected on an Agilent Femto Pulse. DNA was sheared with a Diagenode Megaruptor and size-selected to roughly 25 kb on a BluePippin. A PacBio sequencing library was produced using the SMRTbell Express Template Prep Kit 2.0, and CCS (HiFi) reads were produced on two 8M flow cells. Pacbio HiFi read quality was assessed for read quality versus read distribution (Figure S1) using software *Pauvre* v0.2.3 (Schultz, Ebbert, and De Coster 2019).

The plastid genomes from five *Pyrus* individuals (Table S3) were assembled using *NOVOPlasty* v4.3.1 (Dierckxsens, Mardulyn, and Smits 2016), setting the expected plastid genome size to 130-170 kb and using the seed file provided (https://github.com/ndierckx/NOVOPlasty). The assembled plastid genomes were annotated using *Ge-Seq* v2.0.3 (Tillich et al. 2017) and visualized using *OGDRAW* v1.3.1 (Greiner, Lehwark, and Bock 2019).

### Genome assembly and scaffolding

Raw HiFi reads were assembled into contigs using *hifiasm* v0.16.0 (H. Cheng et al. 2021). To scaffold the “d’Anjou” genome, 1g of young leaf tissue was used as input for a Dovetail Omni-C library per manufacturer instructions (Dovetail Genomics, Inc.). The Omni-C library was sequenced on an Illumina NovaSeq6000 using paired-end 150 base-pair reads. To map the Omni-C data to our preliminary genome assembly, we followed the Arima genomics pipeline (https://github.com/ArimaGenomics/mapping_pipeline). Scaffolding was then performed using yet another Hi-C scaffolding tool (YaHS) with default parameters (Zhou, McCarthy, and Durbin 2023). Omni-C contact maps were visualized using Juicebox version 1.11.08 (Durand et al. 2016). We encountered several examples of likely misassembled regions, which were manually rearranged in Juicebox and documented in Supplementary Methods. We assessed genome completeness using compleasm v0.2.2 with the lineage “embryophyta_odb10” (Huang and Li 2023).

### Annotating repeats and Transposable Elements

Transposable elements (TEs) were predicted and annotated from the pear genome assembly using the Extensive de-novo TE Annotator (EDTA) pipeline (v1.9.3) (Ou et al. 2019; Ellinghaus, Kurtz, and Willhoeft 2008; Xu and Wang 2007; Ou and Jiang 2019, 2018; Su, Gu, and Peterson 2019; Shi and Liang 2019; Xiong et al. 2014). EDTA parameters were set to the following: “--species others --step all --sensitive 1 --anno 1 --evaluate 1 --threads 4”. We calculated the coverage of genes and repeats in 1 Mb windows with a 100 Kb step using bedtools version 2.30.0 (Quinlan and Hall 2010) and plotted these onto the chromosomes using karyoploteR version 1.18.0 (Gel and Serra 2017).

### Structural variant analysis

First, we aligned assemblies using MUMmer (Marçais et al. 2018). Next, we characterized structural variants between genome assemblies using Assemblytics (Nattestad and Schatz 2016). More details are provided in the Supplementary Methods.

### Gene annotation

We annotated protein-coding genes using MAKER2 (Holt and Yandell 2011). Arabidopsis Araport 11 proteins and seven *P. communis* cv. ‘d’Anjou’ RNA-seq libraries were used as evidence (C.-Y. Cheng et al. 2017). RNA-seq libraries are available on the NCBI SRA under Accession PRJNA791346. We performed one round of evidence-based annotation and used that to iteratively train ab-initio prediction models through both SNAP and Augustus. More details are provided in Supplementary Methods.

### RNA-seq analyses

Reads were adapter trimmed using the BBMap ‘bbduk.sh’ script (*BBMap: A Fast, Accurate, Splice-Aware Aligner* 2014). Gene expression was quantified using Kallisto (Bray et al. 2016). More details are provided in the Supplementary Methods.

### Comparative genomic analyses

We identified putative synteny constrained orthologs between *Pyrus communis* cv. ‘d’Anjou’, *Malus domestica* ‘Honeycrisp’, and *Prunus cerasus* ‘Montmorency’ using the JCVI utilities library compara catalog ortholog function (Tang et al. 2015). Synonymous substitution rates were calculated using a custom Ka/Ks pipeline (https://github.com/Aeyocca/ka_ks_pipe). Briefly, orthologs were aligned using MUSCLEv3.8.31 (Edgar 2004), and PAL2NAL v14 was used to convert the peptide alignment to a nucleotide alignment and Ks values were computed between gene pairs using codeml from PAML v4.9 with parameters specified in the control file found in the GitHub repository listed above (Suyama, Torrents, and Bork 2006; Yang 1997).

## Results

### Nuclear Genome assembly

We generated several types of sequencing data to assemble and annotate the d’Anjou genome (Fig 1). Given an estimated genome size of ∼550Mb (Niu et al. 2020), we generated 113X coverage of Illumina shotgun data, 66X coverage of Pacbio HiFi data and 190X of Omni-C data per haplotype. Genomescope estimated a k-mer based genome size of ∼495Mb, 46.79% of repeated sequences, and 1.79% heterozygosity (Fig S1). We assessed the quality of our HiFi reads using Pauvre indicating high quality libraries and a read length distribution centered around 15kb (Fig S2). Our mean and median read lengths were 15,555bp and 14,758bp while the longest read was 49,417bp long.

**Figure 1:**
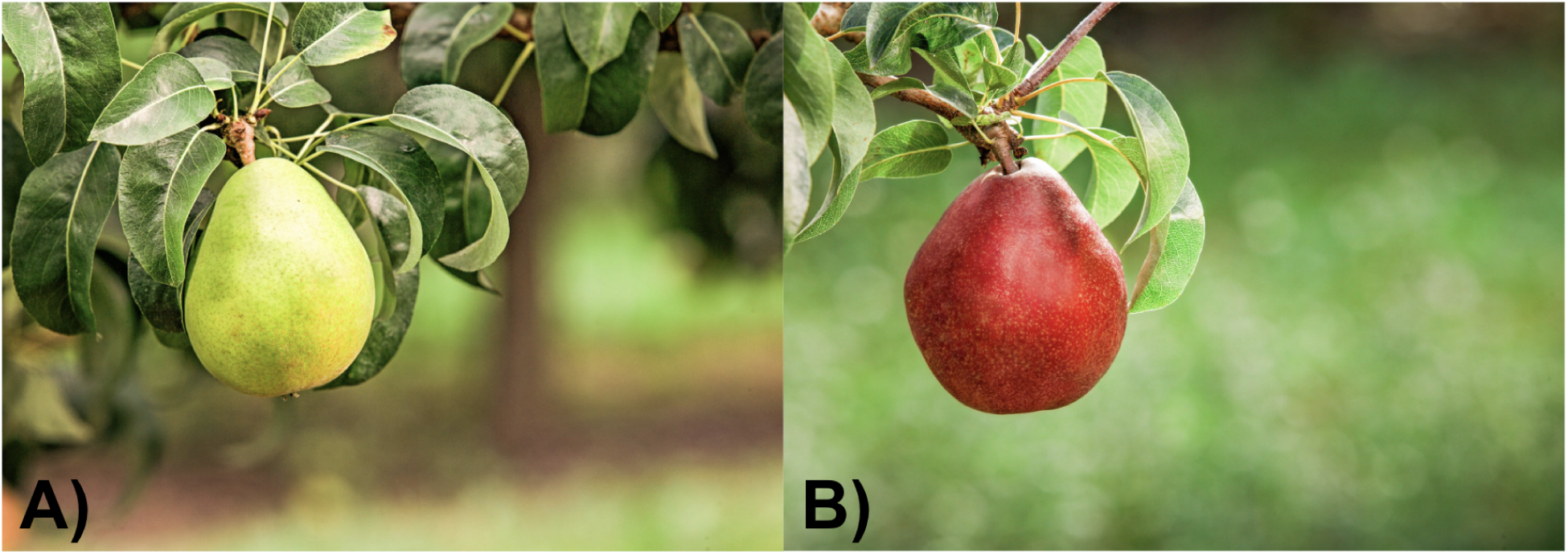
Pear fruit photographs. Photographs of green ‘d’Anjou’ fruit (A) and red ‘d’Anjou’ fruit (B). Photos were provided by USA Pears.

The final assembly is haplotype-resolved with 17 chromosomes per haplotype. Chromosomes were oriented according to the *Malus domestica* “Honeycrisp” assembly (Khan et al. 2022). The final assembly consisted of nearly 540Mb per haplotype with >93% of the raw contig assemblies contained in the 17 chromosomes (Fig S3). The contig N50s for haplotype 1 and 2 respectively were 14.7Mb and 13.4Mb while the scaffold N50s were 29.6Mb. We found >99% complete BUSCOs in each haplotype with over 30% of them present in duplicate, reflecting the whole-genome duplication (WGD) experienced by the Maleae lineage ∼45 million years ago (Xiang et al. 2017). Over 99% of our Illumina reads were properly mapped back to our assembly. Kmer based completeness between Illumina reads and the final assembly demonstrated high quality values (36.16) and low error rates (0.0002423) for both haplotypes.

### Chloroplast assembly

We also assembled the chloroplast of *P. communis* cv. ‘d’Anjou’ along with four other *Pyrus* species or accessions (Table S3; Fig S4; Fig 2). The chloroplast genomes were similar in size, ranging from 159kb to 161kb, and consisted of a large single-copy region, small single-copy region, and two inverted repeats for each species. *Pyrus* as a genus consists of two major genetic groups: Occidental and Oriental (Zheng et al. 2014). *Pyrus hopeiensis*, *P. pyrifolia*, and *P. bretscheirderi* are all considered Oriental species. We estimated phylogenetic relationships between our chloroplast assemblies and found both representatives of *Pyrus communis* sister to each other consistent with expectations.

**Figure 2:**
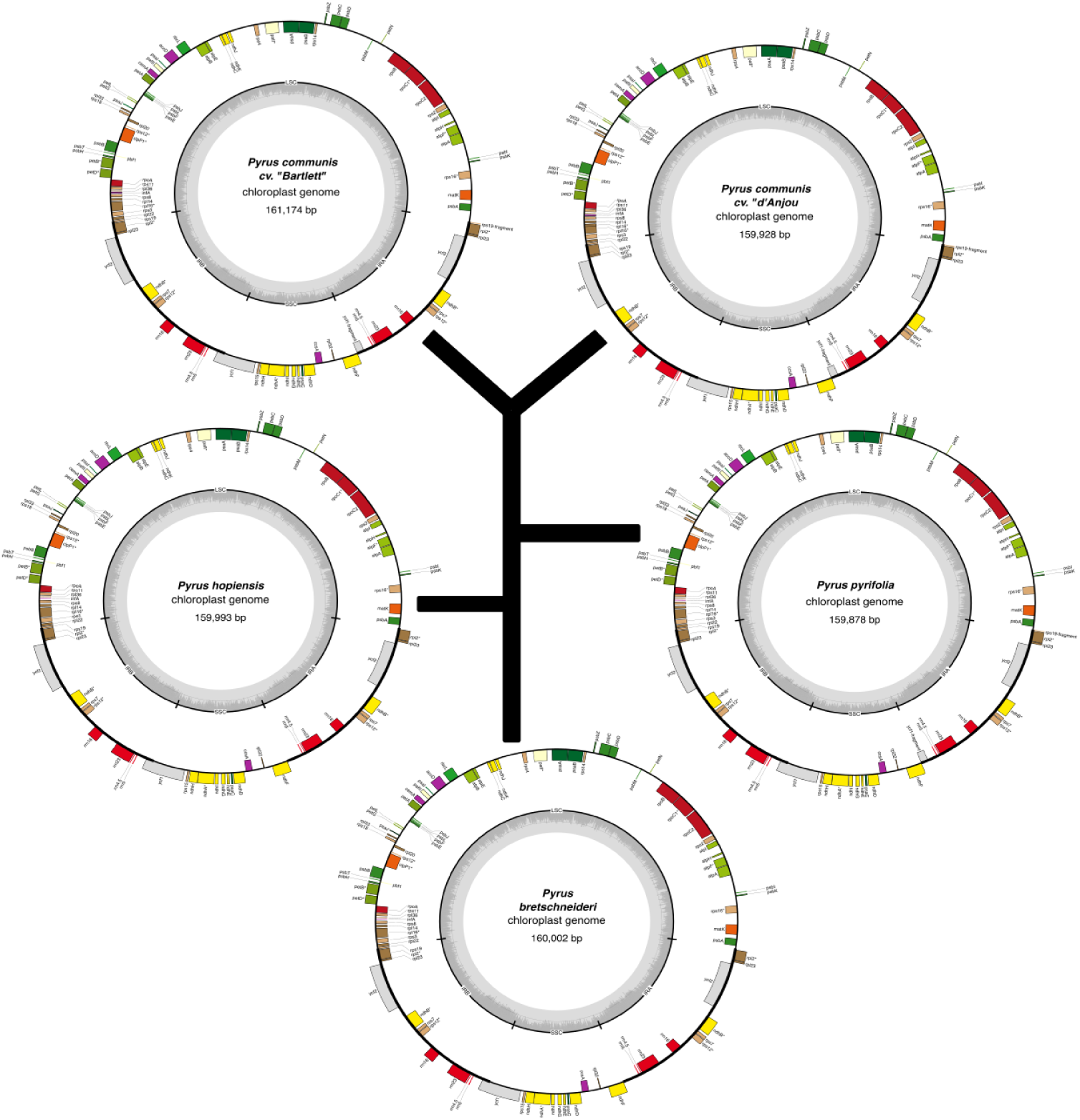
Chloroplast assemblies and phylogeny. Chloroplast genomes of assorted pear cultivars - assemblies and annotations. Plastid assemblies were carried out using *NOVOPlasty* v4.4.1 and annotated using *Ge-Seq* v2.0.3. Phylogenetic relationships were estimated using maximum likelihood under the generalized time reversible model.

Transposable Elements (TEs) are important components of plant genomes, contributing to genome size variation, gene family evolution, and transcriptional novelty (Lu et al. 2019; Quadrana 2020). Repetitive elements were annotated using the Extensive *de novo* Transposable Element Annotator (EDTA; (Ou et al. 2019)) (Table 1). A total of 39-42% of each haplotype consisted of repetitive elements. The majority of these elements by length were long terminal repeat (LTR) retrotransposons accounting for ∼32% of each haplotype. These elements are most abundant around the putative centromeres, but are also ubiquitous in gene rich regions (Fig 3). Terminal inverted repeats (TIRs) were also abundant and dominated by Mutator elements (∼3.4% of each haplotype).

**Figure 3:**
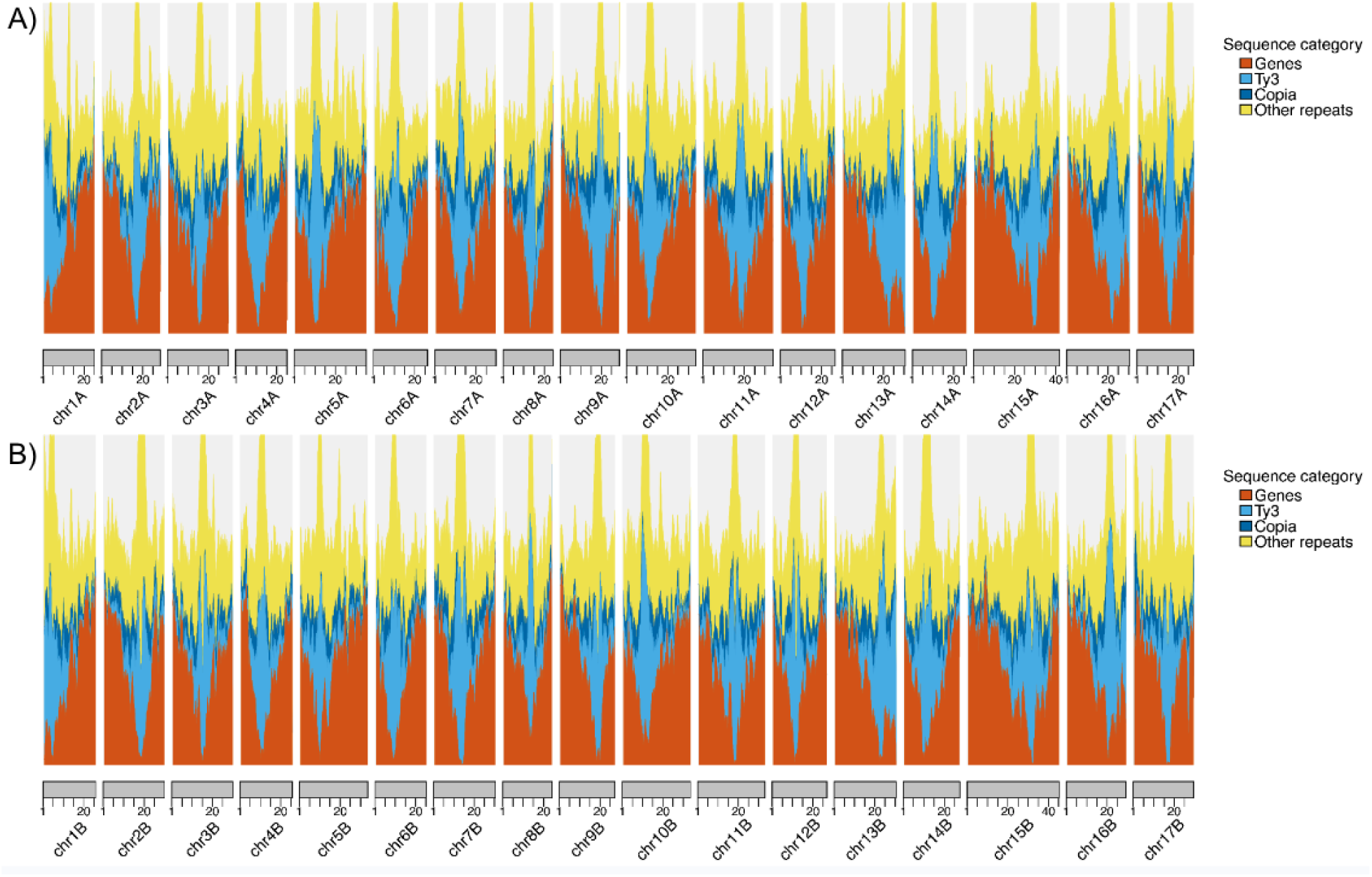
distributions of genomic elements. Density of genomic elements across our assembly. Feature densities are calculated in 1Mb windows with a 100kb step size. Features on haplotype 1 are listed in panel A, and those on haplotype 2 are listed in panel B. Genes are colored orange, Ty3 transposable elements are colored light blue, Copia transposable elements are colored dark blue, and other repeat elements annotated by EDTA are colored yellow. Numbers along the x-axis correspond to position along the chromosome (Mb).

**Table 1:**
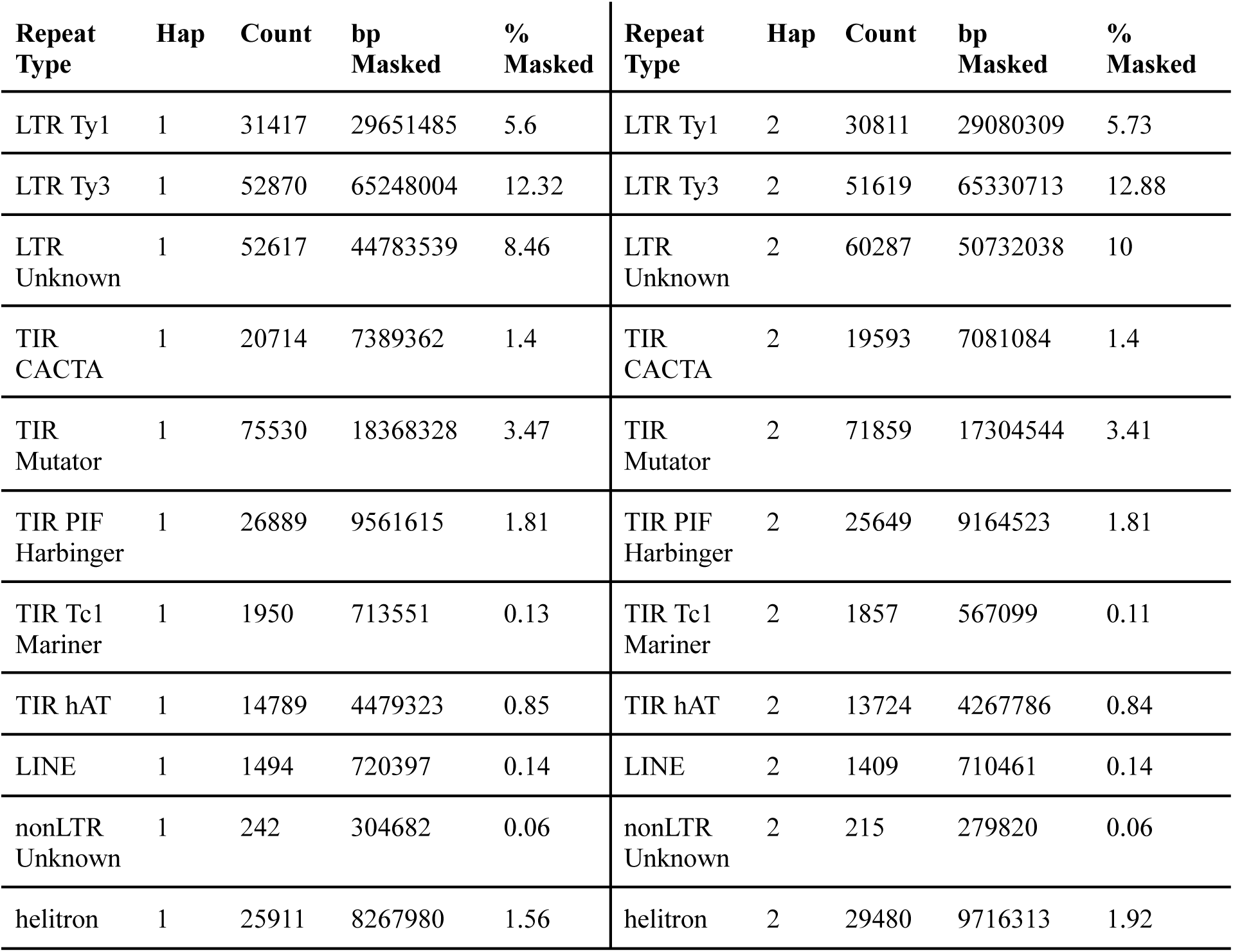
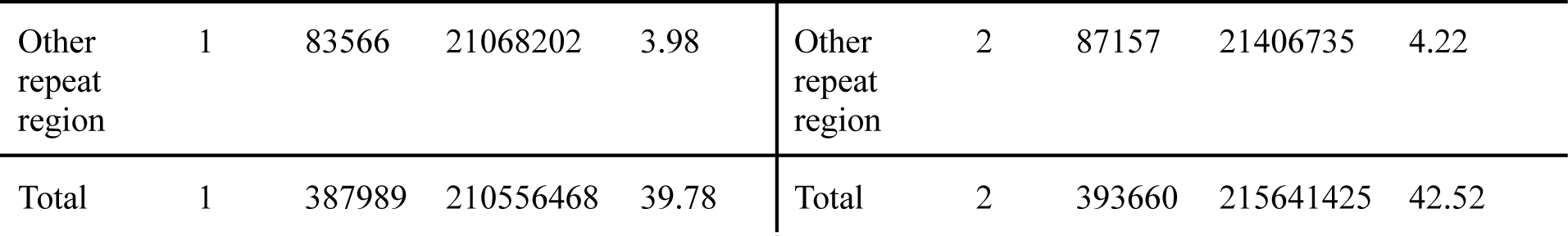
Summary of repeat elements annotated by EDTA. Abbreviations are as follows. LTR; Long-Terminal Repeat. TIR; Terminal Inverted Repeat. PIF; P instability Factor. LINE; Long interspersed nuclear element. Hap; Haplotype. bp; base pairs

Each haplotype was independently annotated with expression evidence, Arabidopsis protein evidence, and *ab initio* gene prediction using the MAKER pipeline (Supplementary Methods; Table S4). We annotated a total of 44,839 genes in haplotype A and 44,561 genes in haplotype B, which is similar to the number of genes annotated in *Malus domestica* ‘Honeycrisp’ (50,105). Gene density was highest on chromosome arms and was inversely related to the density of transposable elements (Fig 3).

There were several structural variants between our two haplotypes (Table 2). We characterized 13,421 variants within 50-10,000 base-pairs between the haplotypes, totaling almost 32Mb of sequence. Repeat expansion and contractions were the largest classes of structural variant. Insertions and deletions also affected nearly 6Mb of sequence between haplotypes. Between *P. communis* cv. ‘d’Anjou’ and *P. communis* cv. ‘Bartlett’, 14,946 variants affected 26Mb of sequence. The total amount of sequence affected is lower than that observed between ‘d’Anjou’ haplotypes. This may simply be due to a more complete assembly for both ‘d’Anjou’ haplotypes relative to the ‘Bartlett’ assembly.

**Table 2:** Structural variants between 50-10,000bp identified by Assemblytics. Indel is short for “Insertion / deletion”.

| Reference | Query | Variant type | # Variants | # bases affected |
| --- | --- | --- | --- | --- |
| ‘d’Anjou’ Hap1 | ‘d’Anjou’ Hap2 | Indel | 4,297 | 6,000,228 |
| ‘d’Anjou’ Hap1 | ‘d’Anjou’ Hap2 | Repeat | 8,711 | 24,943,411 |
| ‘d’Anjou’ Hap1 | ‘Bartlett’ | Indel | 5,739 | 4,439,368 |
| ‘d’Anjou’ Hap1 | ‘Bartlett’ | Repeat | 8,910 | 11,571,098 |

### Comparative genomics and polyploidy

Rosaceae as a plant family contains several important crops such as pear, apple, peach, cherry, and blackberry. Comparative genomics between these crops may allow functional genomics in one species to be translated to others. Therefore, we compared the genomes of three of these important crops: *P. communis* ‘d’Anjou’ (pear), *Malus domestica* ‘Honeycrisp’ (apple), and *Prunus cerasus* ‘Montmorency’ (cherry; (Goeckeritz et al. 2023)). Both our assembled haplotypes were highly collinear with each other and with apple. We identified 40,567 orthologs between pear haplotypes, 30,340 orthologs between pear haplotype 1 and apple, and 20,526 orthologs to *P. cerasus* ‘Montmorency’ consistent with pear’s divergence with apple postdating that to cherry.

Apple and pear share a WGD occurring after their divergence with cherry (Xiang et al. 2017). Our results show they both demonstrate a high percentage (>⅓) of duplicated BUSCO genes as well as 17 chromosomes, almost double the Amygdaloideae base chromosome count of 9 (Hodel et al. 2021). Therefore, we infer apple and pear retain much of their genome in duplicate. Across all genes within *P. communis* cv. ‘d’Anjou’, approximately 57% are classified as having a syntenic paralog retained from this WGD event (Table S5).

‘Montmorency’ is a tetraploid formed from a hybridization between different *Prunus* species after their divergence with the common ancestor of apple and pear. Therefore, we only compared the “A” subgenome to our assemblies. As expected, each cherry “A” subgenome scaffold was syntenic with ∼2 pear and apple scaffolds (Fig 4A). Additionally there were blocks in pear syntenic with 2 regions of apple that are likely regions retained from the last WGD event. There were likely further karyotype changes since the divergence of Malineae and cherry as the syntenic blocks are not entirely retained nor perfectly paired in 1:2 ratios. However, there remains high collinearity with these genomes suggesting future translation of functional genomics across species.

**Figure 4:**
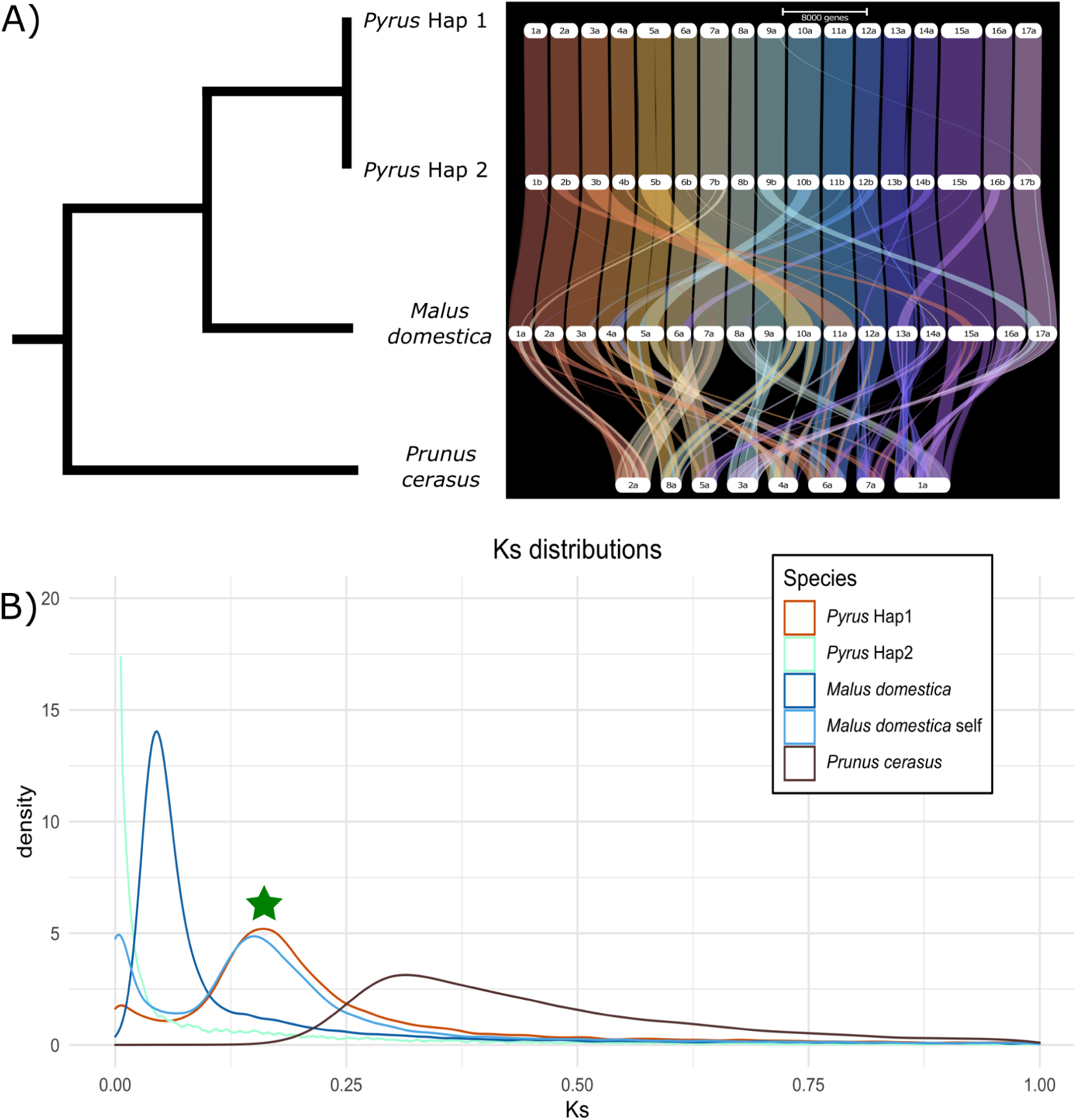
Ribbon plot and Ks distributions. (A) A phylogenetic tree with known relationships between four assemblies. To the right is a ribbon plot based on gene synteny created with GENESPACE (Lovell et al. 2022). (B) A density plot showing the distribution of synonymous substitution rates (Ks) between genome-wide gene pairs. The shared WGD event is denoted by a green star. All comparisons are to *Pyrus communis* cv. ‘d’Anjou’ haplotype 1 except for the “*Malus domestica* self” comparison. Abbreviations are as follows: “*Pyrus* Hap1” - “*Pyrus communis* cv. ‘d’Anjou’ haplotype 1”, “*Pyrus* Hap2” - “*Pyrus communis* cv. ‘d’Anjou’ haplotype 2”.

The distribution of synonymous substitution rates (Ks) across gene pairs indicates the divergence between them as gene pairs will accumulate synonymous substitutions over time (Yang and Nielsen 2000; Senchina et al. 2003). We see orthologs between haplotype 1 and 2 in our assembly have a Ks distribution centered near zero as expected for allelic copies of genes that are still segregating within the species. Comparing haplotype 1 to itself identifies gene pairs that are retained from the most recent WGD event. We see this distribution is higher than that of gene pairs between *Pyrus* and *Malus* suggesting this WGD event occurred before the divergence of these species. Additionally, comparing *M. domestica* to itself shows a distribution similar to that of the *Pyrus* self comparison as expected reflecting a shared WGD event or at the very least, a different WGD event occurring around the same time (Fig 4B; green star). This distribution is lower than that compared to *Prunus cerasus* as this WGD event post-dates the divergence of the cherry and apple/pear lineages.

### Gene expression

We quantified gene expression across seven tissues (Table 3). We found expression evidence for ∼33-35,000 gene models per tissue. Most gene models were expressed in Fruitlet Stage 1, and the least were expressed in Fruitlet Stage 2 suggesting dynamic gene expression across fruit development. There was evidence of gene expression in at least a single tissue for 40,734 gene models, while 2,152 genes were expressed in only a single tissue (average of 307 genes per tissue). Our expression data were generated to assist genome annotation and are only single replicates. We therefore cannot perform differential expression analyses. We instead performed hierarchical clustering of gene expression (Fig 5). We see stable clustering across haplotypes and find similar tissues cluster together. For example, our two fruit libraries clustered with each other. We generated an upset plot showing the fifteen largest intersects of genes expressed >1 transcript per million (TPM; Fig 5). The largest intersect was genes expressed >1 TPM in every tissue queried. The top fifteen intersects, however, included each of the seven tissue-specific categories. Open Buds had the most tissue-specific genes (445) while Budding Leaves specific genes had the least (171).

**Figure 5:**
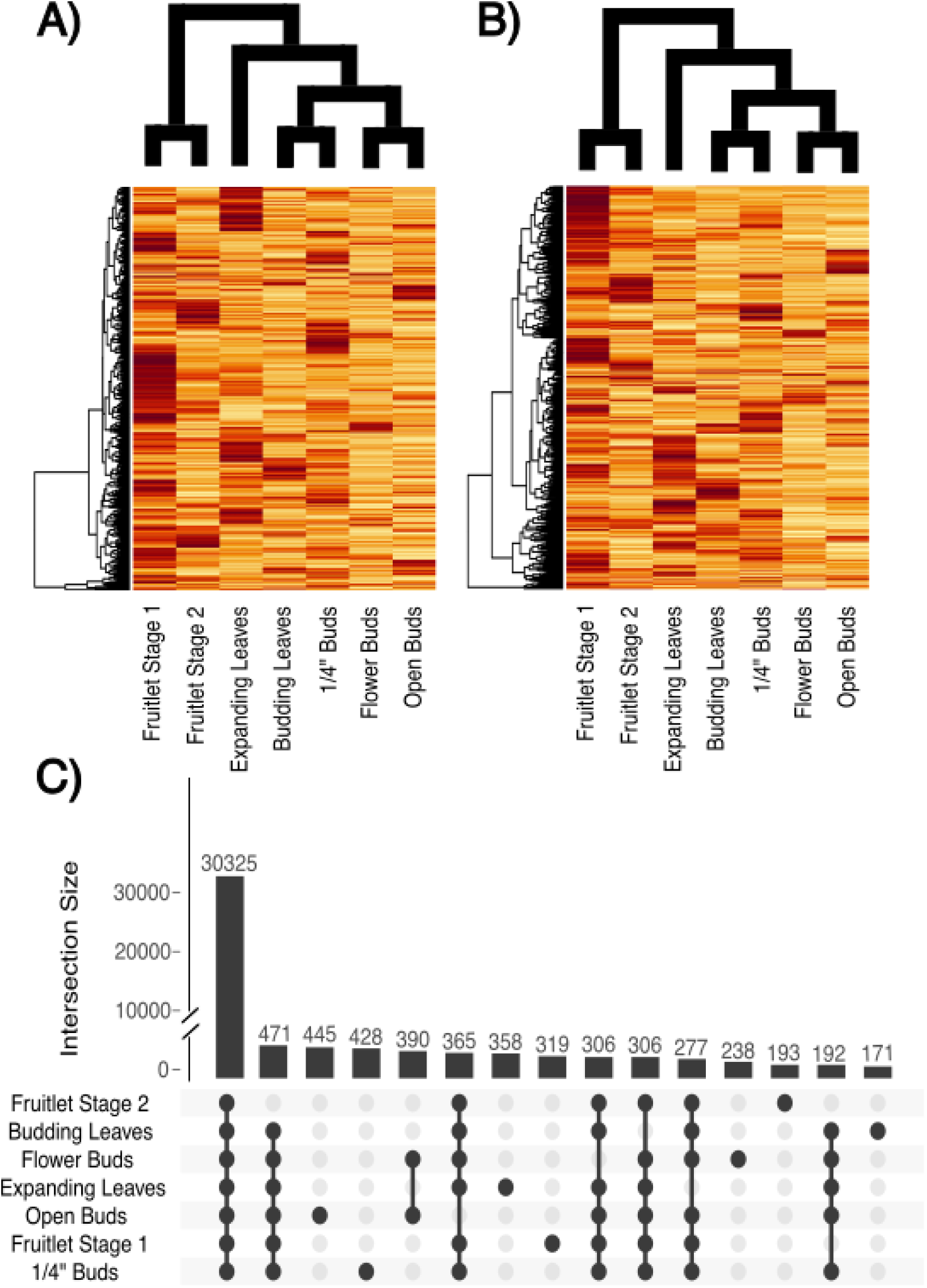
Gene expression characterization. Heatmaps and Upset plot of gene expression. Cladograms represent the relationships between libraries through hierarchical clustering. 1000 genes are displayed that show expression in each tissue and have the highest expression variance. A) represents haplotype 1 and B) represents haplotype 2. C) Upset plot of expression across tissues for haplotype 1. Genes were considered expressed if they had a TPM value above 1. Note the break in the y-axis.

**Table 3:** Expression characteristics of *Pyrus communis* cv. ‘d’Anjou’. Abbreviations are as follows: Hap; Haplotype. TPM; transcripts per million reads.

| Tissue | Hap | Genes expressed | Median TPM | Tissue | Hap | Genes expressed | Median TPM |
| --- | --- | --- | --- | --- | --- | --- | --- |
| Budding Leaves | 1 | 33594 | 84.97 | Budding Leaves | 2 | 33470 | 88.00 |
| Expanding Leaves | 1 | 34469 | 119.7 | Expanding Leaves | 2 | 34380 | 122.0 |
| Flower Buds | 1 | 34138 | 71.34 | Flower Buds | 2 | 34082 | 73.3 |
| Fruitlet Stage 1 | 1 | 34923 | 193 | Fruitlet Stage 1 | 2 | 34797 | 200 |
| Fruitlet Stage 2 | 1 | 33227 | 96.4 | Fruitlet Stage 2 | 2 | 33107 | 100.0 |
| Open Buds | 1 | 34463 | 72.0 | Open Buds | 2 | 34372 | 74.02 |
| ¼” buds | 1 | 34718 | 108.3 | ¼” buds | 2 | 34513 | 111.00 |

## Conclusion

We assembled a chromosome-scale phased genome assembly for cultivated European pear. PacBio HiFi reads coupled with Dovetail Omni-C resulted in a high quality assembly, displaying high kmer completeness, quality scores, synteny with available assemblies, and recovery of universal single-copy orthologs. This assembly revealed thousands of structural variants between haplotypes which are of great importance to future pear breeding efforts as structural variants disrupt recombination. Comparative analyses between other members of the Rosaceae family demonstrated deeply conserved synteny and recovered evidence for a 45 million year old whole genome duplication event. Gene expression across several tissue types was largely conserved, but thousands of genes also constrained themselves to a single tissue.

Further characterization of pear germplasm will accelerate breeding gains not only within pear but potentially across multiple Rosaceous crops. Lastly, we highlight the utility of generating such genomes as part of semester courses, and the training opportunities that it provides.

## Data Availability

Data used to generate this assembly are deposited in the NCBI SRA under BioProject PRJNA992953. Gene expression data are available separately under BioProject PRJNA791346. Custom scripts used throughout are available on github https://github.com/Aeyocca/dAnjou_genome_MS.

## Acknowledgements

This work was supported through funding from NSF PGRP CAREER award #2239530 to AH, NSF IOS-EDGE award #2128196 to AH, Washington Tree Fruit Research Commission award AP-19-103 to LH, SF, AH, the USDA ARS, and the Auburn University Department of Crop, Soil, and Environmental Sciences who supported student compute costs for the CSES7120 Plant Genomics course.

---

**Figure S1:**
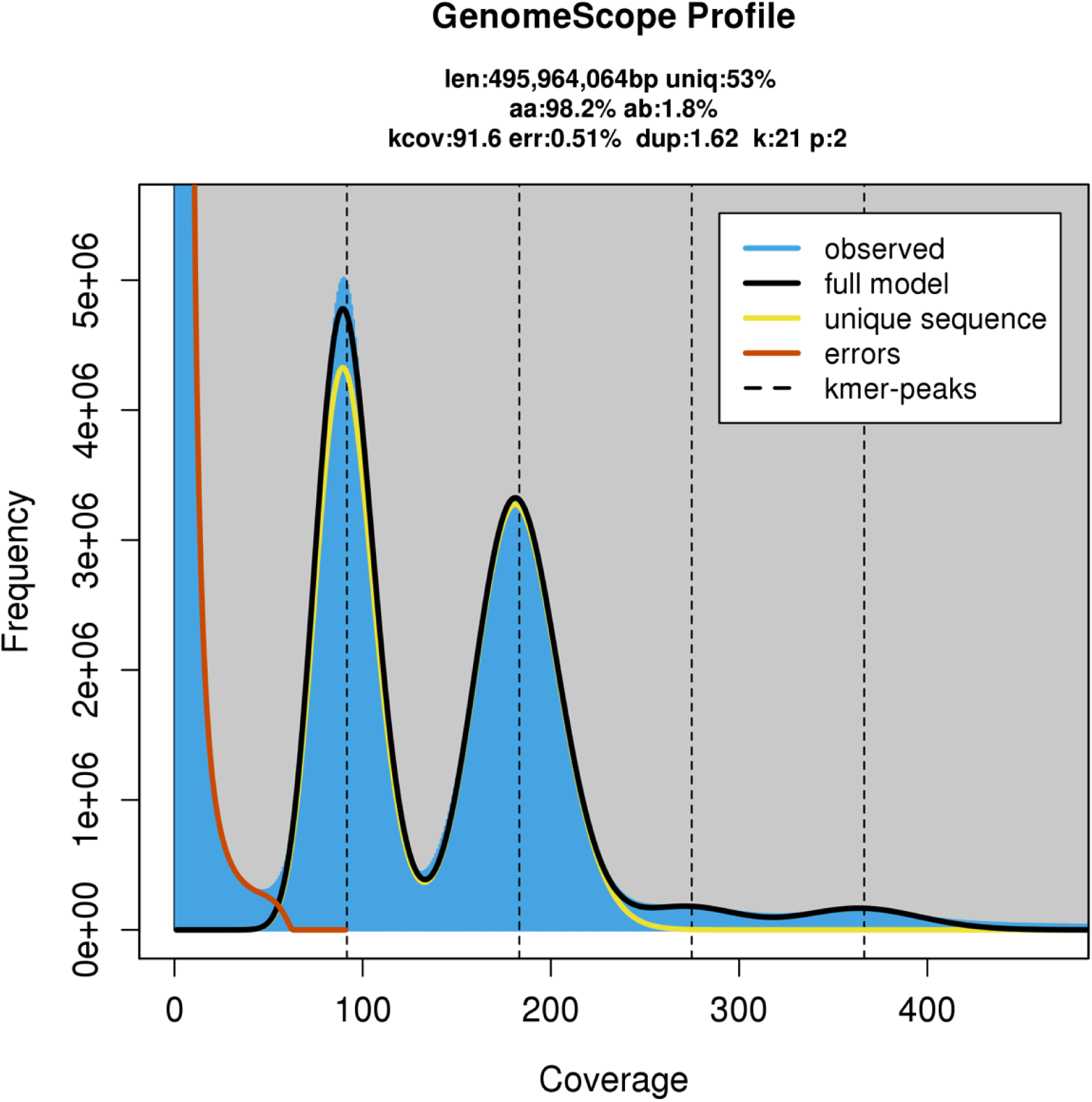
GenomeScope output for ‘d’Anjou’ short-read data. GenomeScope k-mer (k=21) profile plot showing there are two major diploid peaks. The tall peak at 91.6X coverage indicates high heterozygosity in this genome.

**Figure S2:**
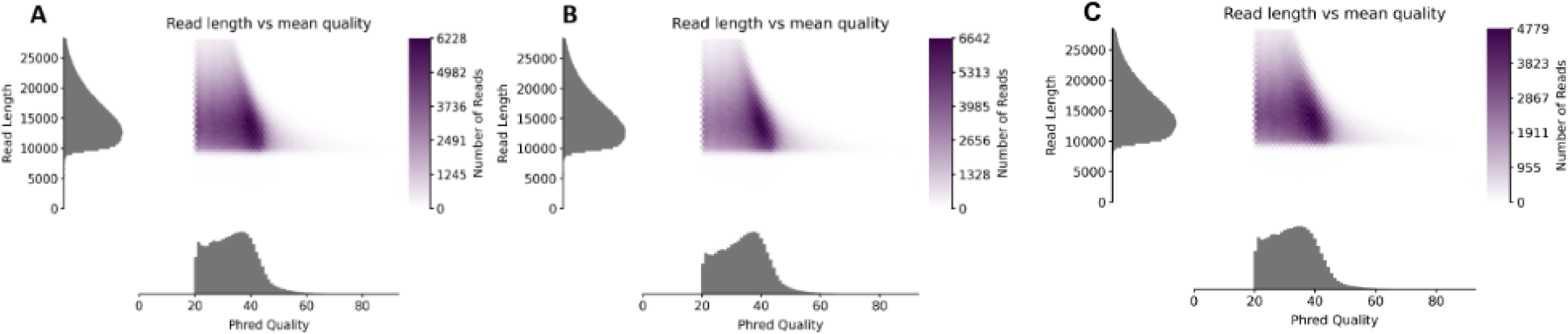
Phred quality analysis of three Pacbio HiFi cells; (A) m64017_211217_051206, (B) m64017_211218_161111, and (C) m64017_211210_084732

**Figure S3:**
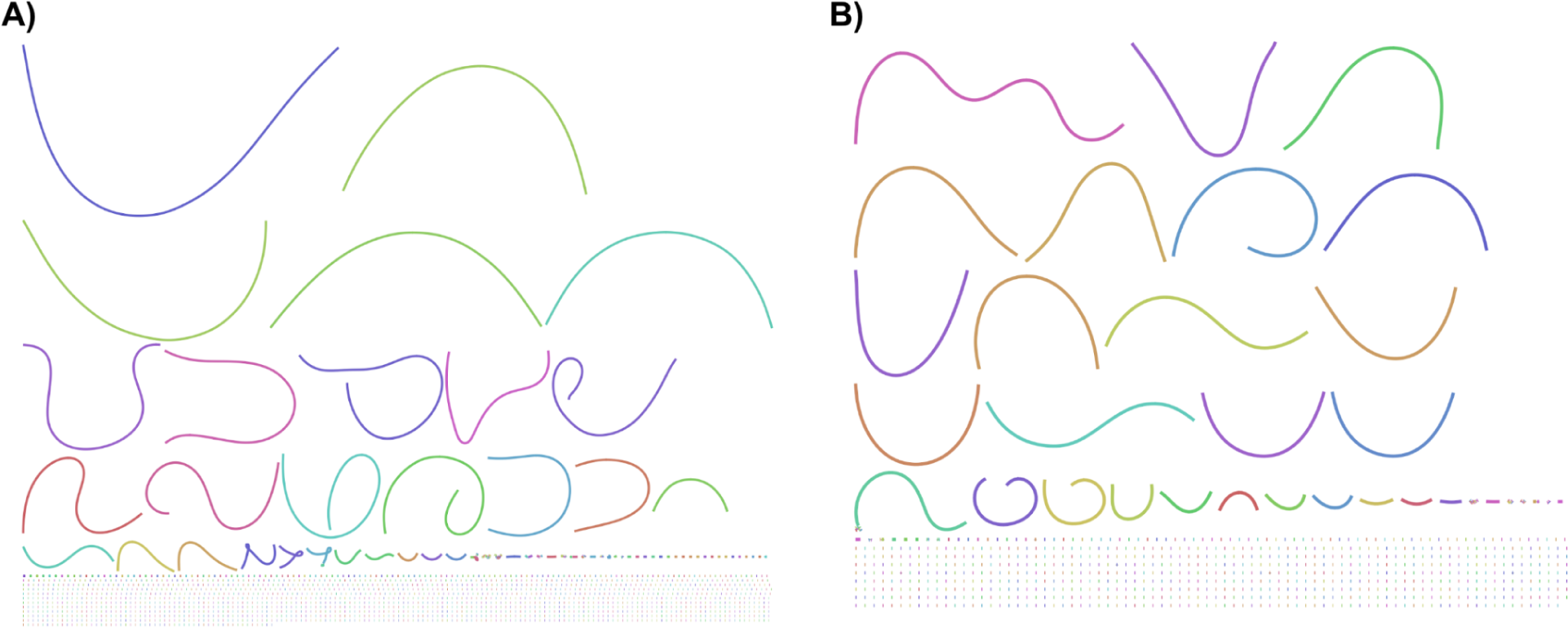
Bandage plots (Wick et al. 2015) for haplotype 1 (A) and haplotype 2 (B).

**Figure S4:**
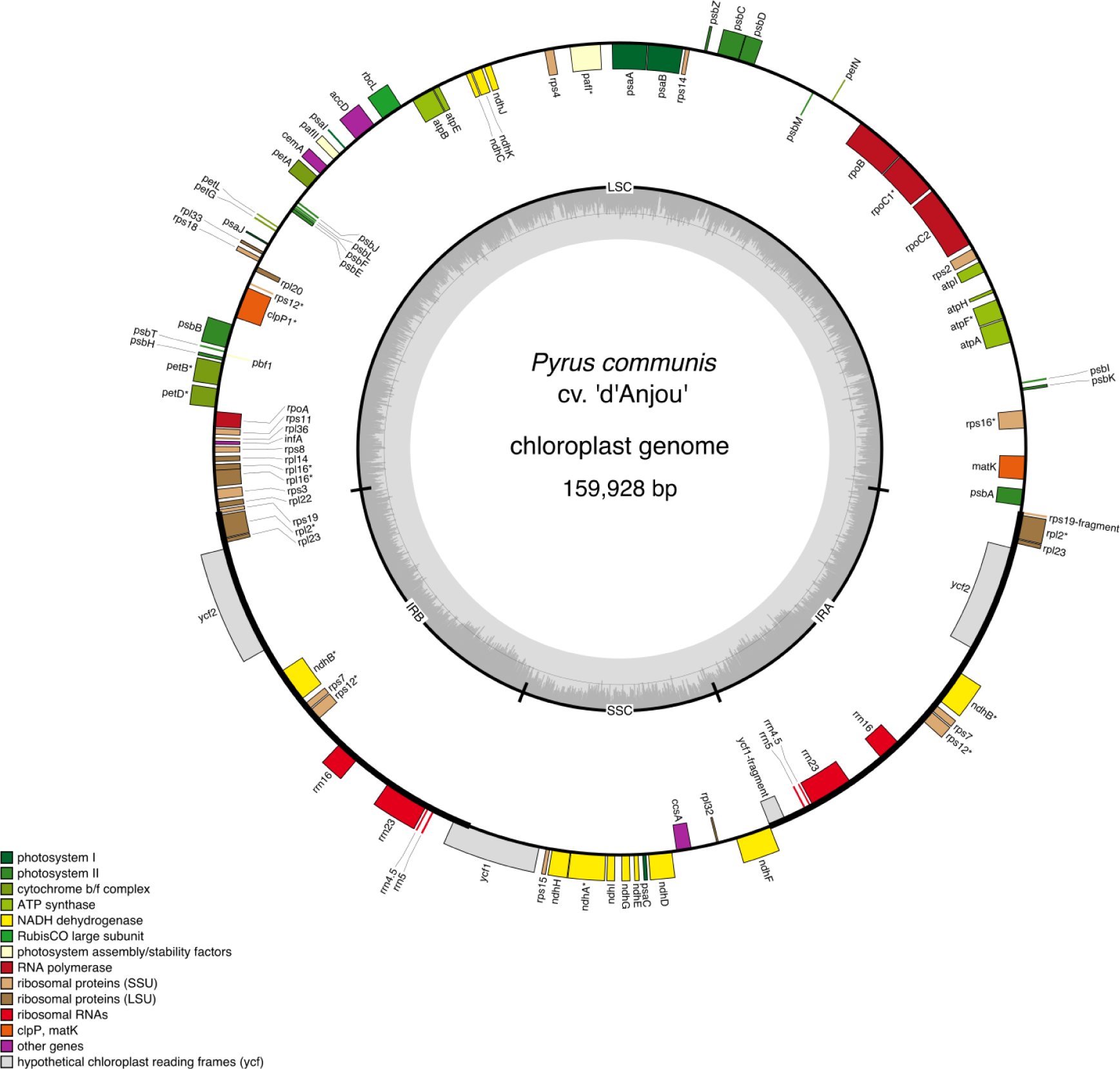
The ‘d’Anjou’ pear chloroplast genome assembly and annotation. Plastid assembly was carried out using *NOVOPlasty* v4.4.1 and annotated using *Ge-Seq* v2.0.3.

**Table S1:**
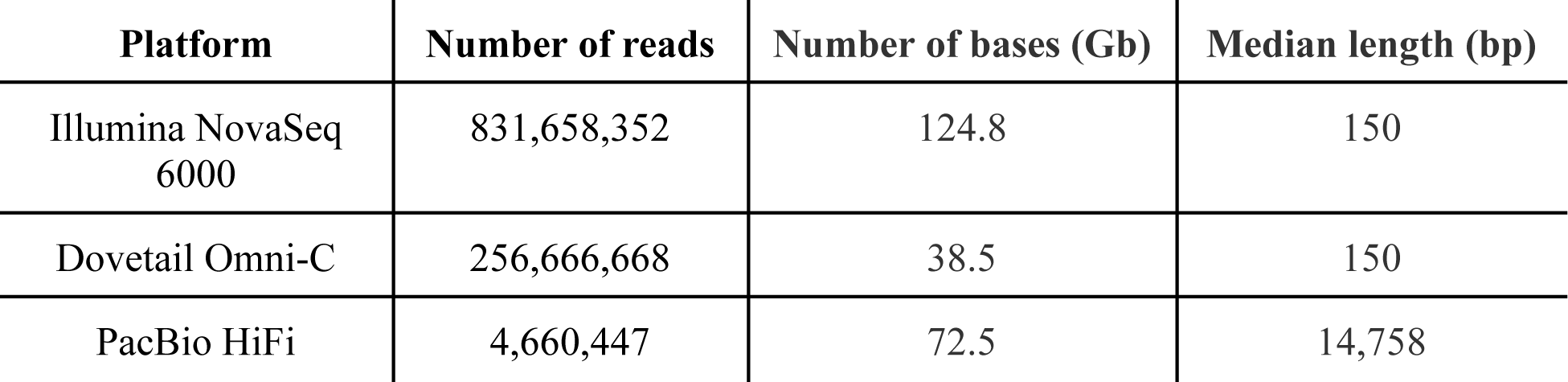
Summary of sequencing reads used for genome assembly.

| Platform | Number of reads | Number of bases (Gb) | Median length (bp) |
| --- | --- | --- | --- |
| Illumina NovaSeq 6000 | 831,658,352 | 124.8 | 150 |
| Dovetail Omni-C | 256,666,668 | 38.5 | 150 |
| PacBio HiFi | 4,660,447 | 72.5 | 14,758 |

**Table S2:**
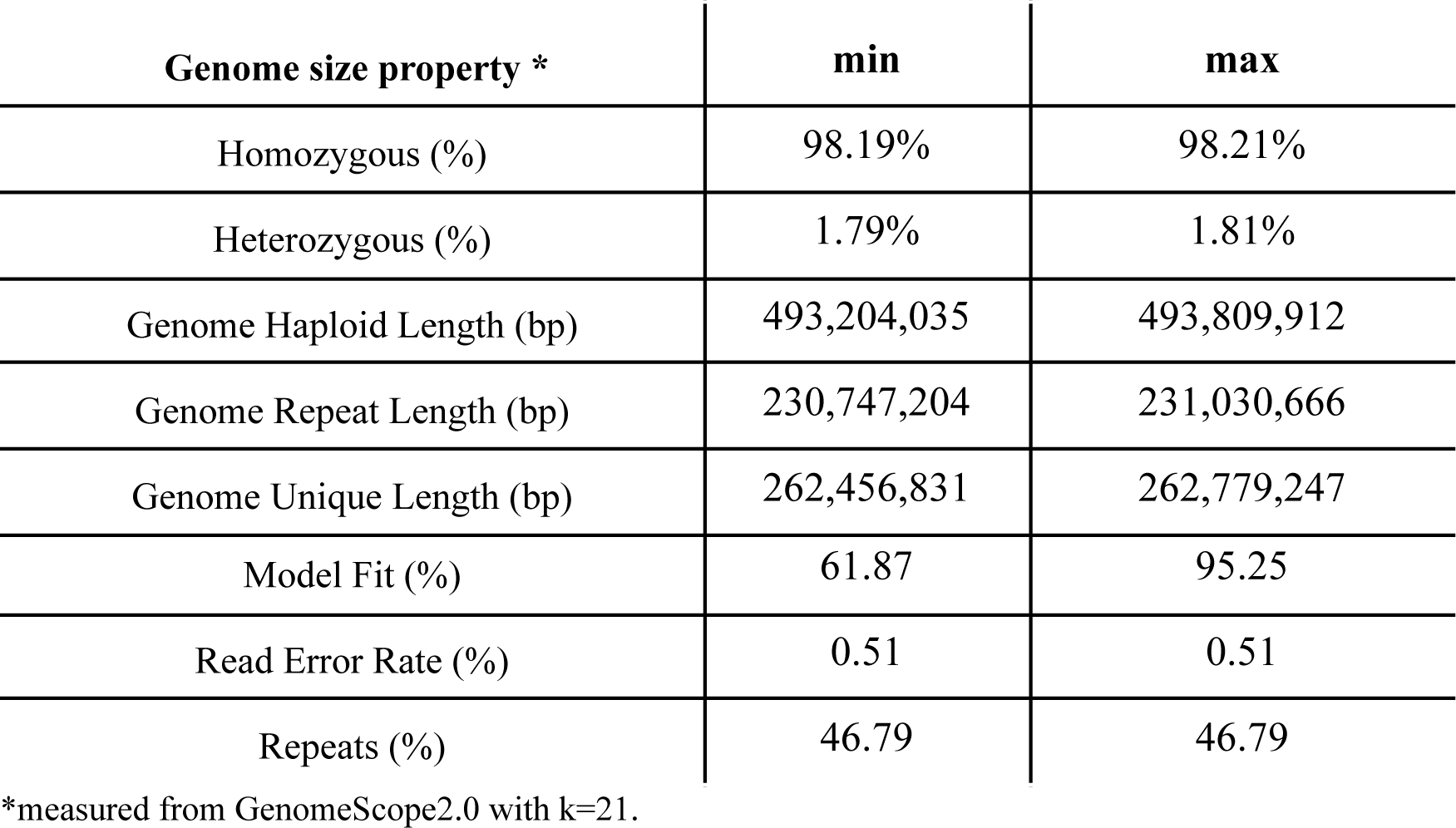
Characteristics of ‘d’Anjou’ pear based on the short-read kmer profile.

| Genome size property * | min | max |
| --- | --- | --- |
| Homozygous (%) | 98.19% | 98.21% |
| Heterozygous (%) | 1.79% | 1.81% |
| Genome Haploid Length (bp) | 493,204,035 | 493,809,912 |
| Genome Repeat Length (bp) | 230,747,204 | 231,030,666 |
| Genome Unique Length (bp) | 262,456,831 | 262,779,247 |
| Model Fit (%) | 61.87 | 95.25 |
| Read Error Rate (%) | 0.51 | 0.51 |
| Repeats (%) | 46.79 | 46.79 |

\*measured from GenomeScope2.0 with k=21.

**Table S3:** Source of data for chloroplast assemblies.

| Species | Source | Number of bases (Gb) | SRAID | # of assembled contigs | # of base-pairs |
| --- | --- | --- | --- | --- | --- |
| *Pyrus hopeiensis* HB-1 | (Y. Li et al. 2021) | 16.34 | SRR14318823 | 3 | 159993 |
| *Pyrus pyrifolia* | (M.-Y. Zhang et al. 2021) | 50.04 | DRR385062 | 4 | 159875 |
| *Pyrus communis* “Bartlett DH” | (Linsmith et al. 2019) | 29.27 | SRR10030340 | 5 | 161148 |
| *Pyrus x bretschneideri* | (Y. Li et al. 2021) | 21.65 | SRR14318827 | 3 | 160001 |
| *Pyrus communis* cv. “d’Anjou” | This publication |  | NA | 1 | 159928 |

**Table S4:** Annotation statistics. Abbreviations are as follows: C (complete), S (single-copy), D (duplicated), F (fragmented), M (missing), n (number).

| Haplotype | # Genes | # Transcripts | Annotated bases | BUSCO string |
| --- | --- | --- | --- | --- |
| 1 | 44558 | 44839 | 68,452,305 | C:97.1%[S:63.1%,D:34.0%],F:1.3%,M:1.6%,n:1614 |
| 2 | 44349 | 44561 | 68,084,531 | C:97.6%[S:64.3%,D:33.3%],F:0.9%,M:1.5%,n:1614 |

**Table S5:** Duplication classification. WGD stands for “Whole Genome Duplication”.

| Duplication type | Haplotype | # of genes | % of genes |
| --- | --- | --- | --- |
| Singleton | 1 | 5726 | 12.77 |
| Dispersed | 1 | 6810 | 15.19 |
| Proximal | 1 | 2865 | 6.39 |
| Tandem | 1 | 3844 | 8.57 |
| WGD or segmental | 1 | 25594 | 57.08 |
| Singleton | 2 | 5818 | 13.06 |
| Dispersed | 2 | 6780 | 15.21 |
| Proximal | 2 | 2839 | 6.37 |
| Tandem | 2 | 4041 | 9.07 |
| WGD or segmental | 2 | 25083 | 56.29 |

## References

Andrews, S., F. Krueger, A. Segonds-Pichon, L. Biggins, C. Krueger, and S. Wingett. 2010. “FastQC: A Quality Control Tool for High Throughput Sequence Data. Babraham Bioinformatics. 2010.”

BBMap: A Fast, Accurate, Splice-Aware Aligner. 2014.

Bray, Nicolas L., Harold Pimentel, Páll Melsted, and Lior Pachter. 2016. “Near-Optimal Probabilistic RNA-Seq Quantification.” Nature Biotechnology 34 (5): 525–27.

Chagné, David, Ross N. Crowhurst, Massimo Pindo, Amali Thrimawithana, Cecilia Deng, Hilary Ireland, Mark Fiers, et al. 2014. “The Draft Genome Sequence of European Pear (Pyrus Communis L. ‘Bartlett’).” PloS One 9 (4): e92644.

Cheng, Chia-Yi, Vivek Krishnakumar, Agnes P. Chan, Françoise Thibaud-Nissen, Seth Schobel, and Christopher D. Town. 2017. “Araport11: A Complete Reannotation of the Arabidopsis Thaliana Reference Genome.” The Plant Journal: For Cell and Molecular Biology 89 (4): 789–804.

Cheng, Haoyu, Gregory T. Concepcion, Xiaowen Feng, Haowen Zhang, and Heng Li. 2021. “Haplotype-Resolved de Novo Assembly Using Phased Assembly Graphs with Hifiasm.” Nature Methods 18 (2): 170–75.

Dierckxsens, Nicolas, Patrick Mardulyn, and Guillaume Smits. 2016. “NOVOPlasty: De Novo Assembly of Organelle Genomes from Whole Genome Data.” Nucleic Acids Research 45 (4): e18–e18.

Doyle, J. J., and J. L. Doyle. 1987. “A Rapid DNA Isolation Procedure for Small Quantities of Fresh Leaf Tissue.” RESEARCH. https://worldveg.tind.io/record/33886/.

Durand, Neva C., James T. Robinson, Muhammad S. Shamim, Ido Machol, Jill P. Mesirov, Eric S. Lander, and Erez Lieberman Aiden. 2016. “Juicebox Provides a Visualization System for Hi-C Contact Maps with Unlimited Zoom.” Cell Systems 3 (1): 99–101.

Edgar, Robert C. 2004. “MUSCLE: Multiple Sequence Alignment with High Accuracy and High Throughput.” Nucleic Acids Research 32 (5): 1792–97.

Ellinghaus, David, Stefan Kurtz, and Ute Willhoeft. 2008. “LTRharvest, an Efficient and Flexible Software for de Novo Detection of LTR Retrotransposons.” BMC Bioinformatics 9 (January): 18.

Ewels, Philip, Måns Magnusson, Sverker Lundin, and Max Käller. 2016. “MultiQC: Summarize Analysis Results for Multiple Tools and Samples in a Single Report.” Bioinformatics 32 (19): 3047–48.

Gel, Bernat, and Eduard Serra. 2017. “karyoploteR: An R/Bioconductor Package to Plot Customizable Genomes Displaying Arbitrary Data.” Bioinformatics 33 (19): 3088–90.

Goeckeritz, Charity Z., Kathleen E. Rhoades, Kevin L. Childs, Amy F. Iezzoni, Robert VanBuren, and Courtney A. Hollender. 2023. “Genome of Tetraploid Sour Cherry (Prunus Cerasus L.) ‘Montmorency’ Identifies Three Distinct Ancestral Prunus Genomes.” Horticulture Research 10 (7). 10.1093/hr/uhad097.

Greiner, Stephan, Pascal Lehwark, and Ralph Bock. 2019. “OrganellarGenomeDRAW (OGDRAW) Version 1.3.1: Expanded Toolkit for the Graphical Visualization of Organellar Genomes.” Nucleic Acids Research 47 (W1): W59–64.

Hodel, Richard G. J., Elizabeth A. Zimmer, Bin-Bin Liu, and Jun Wen. 2021. “Synthesis of Nuclear and Chloroplast Data Combined With Network Analyses Supports the Polyploid Origin of the Apple Tribe and the Hybrid Origin of the Maleae-Gillenieae Clade.” Frontiers in Plant Science 12: 820997.

Holt, Carson, and Mark Yandell. 2011. “MAKER2: An Annotation Pipeline and Genome-Database Management Tool for Second-Generation Genome Projects.” BMC Bioinformatics 12 (December): 491.

Huang, Neng, and Heng Li. 2023. “miniBUSCO: A Faster and More Accurate Reimplementation of BUSCO.” *bioRxiv*, June. 10.1101/2023.06.03.543588.

Khan, Awais, Sarah B. Carey, Alicia Serrano, Huiting Zhang, Heidi Hargarten, Haley Hale, Alex Harkess, and Loren Honaas. 2022. “A Phased, Chromosome-Scale Genome of ‘Honeycrisp’ Apple ().” GigaByte (Hong Kong, China) 2022 (September): gigabyte69.

Li, Jiaming, Mingyue Zhang, Xiaolong Li, Awais Khan, Satish Kumar, Andrew Charles Allan, Kui Lin-Wang, et al. 2022. “Pear Genetics: Recent Advances, New Prospects, and a Roadmap for the Future.” Horticulture Research 9 (January). 10.1093/hr/uhab040.

Linsmith, Gareth, Stephane Rombauts, Sara Montanari, Cecilia H. Deng, Jean-Marc Celton, Philippe Guérif, Chang Liu, et al. 2019. “Pseudo-Chromosome–length Genome Assembly of a Double Haploid ‘Bartlett’ Pear (Pyrus Communis L.).” GigaScience 8 (12): giz138.

Li, Yongtan, Jun Zhang, Shijie Wang, Yiwen Zhang, and Minsheng Yang. 2021. “The Distribution and Origins of Pyrus Hopeiensis-‘Wild Plant With Tiny Population’ Using Whole Genome Resequencing.” Frontiers in Plant Science 12 (June): 668796.

Lovell, John T., Avinash Sreedasyam, M. Eric Schranz, Melissa Wilson, Joseph W. Carlson, Alex Harkess, David Emms, David M. Goodstein, and Jeremy Schmutz. 2022. “GENESPACE Tracks Regions of Interest and Gene Copy Number Variation across Multiple Genomes.” eLife 11 (September). 10.7554/eLife.78526.

Lu, Zefu, Alexandre P. Marand, William A. Ricci, Christina L. Ethridge, Xiaoyu Zhang, and Robert J. Schmitz. 2019. “The Prevalence, Evolution and Chromatin Signatures of Plant Regulatory Elements.” Nature Plants 5 (12): 1250–59.

Marçais, Guillaume, Arthur L. Delcher, Adam M. Phillippy, Rachel Coston, Steven L. Salzberg, and Aleksey Zimin. 2018. “MUMmer4: A Fast and Versatile Genome Alignment System.” PLoS Computational Biology 14 (1): e1005944.

Marçais, Guillaume, and Carl Kingsford. 2011. “A Fast, Lock-Free Approach for Efficient Parallel Counting of Occurrences of K-Mers.” Bioinformatics 27 (6): 764–70.

Nattestad, Maria, and Michael C. Schatz. 2016. “Assemblytics: A Web Analytics Tool for the Detection of Variants from an Assembly.” Bioinformatics 32 (19): 3021–23.

Niu, Yingying, Weiquan Zhou, Xiangying Chen, Guoquan Fan, Shikui Zhang, and Kang Liao. 2020. “Genome Size and Chromosome Ploidy Identification in Pear Germplasm Represented by Asian Pears - Local Pear Varieties.” Scientia Horticulturae. 10.1016/j.scienta.2020.109202.

Ou, Shujun, and Ning Jiang. 2018. “LTR_retriever: A Highly Accurate and Sensitive Program for Identification of Long Terminal Repeat Retrotransposons.” Plant Physiology 176 (2): 1410–22.

Ou, Shujun, and Ning Jiang. 2019 “LTR_FINDER_parallel: Parallelization of LTR_FINDER Enabling Rapid Identification of Long Terminal Repeat Retrotransposons.” Mobile DNA 10 (December): 48.

Ou, Shujun, Weija Su, Yi Liao, Kapeel Chougule, Jireh R. A. Agda, Adam J. Hellinga, Carlos Santiago Blanco Lugo, et al. 2019. “Benchmarking Transposable Element Annotation Methods for Creation of a Streamlined, Comprehensive Pipeline.” Genome Biology 20 (1): 275.

Quadrana, Leandro. 2020. “The Contribution of Transposable Elements to Transcriptional Novelty in Plants: The Affair.” Transcription 11 (3-4): 192–98.

Quinlan, Aaron R., and Ira M. Hall. 2010. “BEDTools: A Flexible Suite of Utilities for Comparing Genomic Features.” Bioinformatics 26 (6): 841–42.

Ranallo-Benavidez, T. Rhyker, Kamil S. Jaron, and Michael C. Schatz. 2020. “GenomeScope 2.0 and Smudgeplot for Reference-Free Profiling of Polyploid Genomes.” Nature Communications 11 (1): 1432.

Schultz, Darrin, Mark Ebbert, and Wouter De Coster. 2019. “Pauvre.” May 16, 2019. https://github.com/conchoecia/pauvre.

Senchina, David S., Ines Alvarez, Richard C. Cronn, Bao Liu, Junkang Rong, Richard D. Noyes, Andrew H. Paterson, Rod A. Wing, Thea A. Wilkins, and Jonathan F. Wendel. 2003. “Rate Variation among Nuclear Genes and the Age of Polyploidy in Gossypium.” Molecular Biology and Evolution 20 (4): 633–43.

Shi, Jieming, and Chun Liang. 2019. “Generic Repeat Finder: A High-Sensitivity Tool for Genome-Wide De Novo Repeat Detection.” Plant Physiology 180 (4): 1803–15.

Su, Weijia, Xun Gu, and Thomas Peterson. 2019. “TIR-Learner, a New Ensemble Method for TIR Transposable Element Annotation, Provides Evidence for Abundant New Transposable Elements in the Maize Genome.” Molecular Plant 12 (3): 447–60.

Suyama, Mikita, David Torrents, and Peer Bork. 2006. “PAL2NAL: Robust Conversion of Protein Sequence Alignments into the Corresponding Codon Alignments.” Nucleic Acids Research 34 (Web Server issue): W609–12.

Tang, H., V. Krishnakumar, J. Li, and X. Zhang. 2015. “Jcvi: JCVI Utility Libraries.” Zenodo. 10.5281/zenodo.31631.

Tillich, Michael, Pascal Lehwark, Tommaso Pellizzer, Elena S. Ulbricht-Jones, Axel Fischer, Ralph Bock, and Stephan Greiner. 2017. “GeSeq - Versatile and Accurate Annotation of Organelle Genomes.” Nucleic Acids Research 45 (W1): W6–11.

United States Department of Agriculture National Agricultural Statistics Service. 2023. “Noncitrus Fruits and Nuts 2022 Summary,” May. https://downloads.usda.library.cornell.edu/usda-esmis/files/zs25x846c/zk51wx21m/k356bk 214/ncit0523.pdf.

Wick, Ryan R., Mark B. Schultz, Justin Zobel, and Kathryn E. Holt. 2015. “Bandage: Interactive Visualization of de Novo Genome Assemblies.” Bioinformatics 31 (20): 3350–52.

Wu, Jun, Yingtao Wang, Jiabao Xu, Schuyler S. Korban, Zhangjun Fei, Shutian Tao, Ray Ming, et al. 2018. “Diversification and Independent Domestication of Asian and European Pears.” Genome Biology 19 (1): 77.

Wu, Jun, Zhiwen Wang, Zebin Shi, Shu Zhang, Ray Ming, Shilin Zhu, M. Awais Khan, et al. 2013. “The Genome of the Pear (Pyrus Bretschneideri Rehd.).” Genome Research 23 (2): 396–408.

Xiang, Yezi, Chien-Hsun Huang, Yi Hu, Jun Wen, Shisheng Li, Tingshuang Yi, Hongyi Chen, Jun Xiang, and Hong Ma. 2017. “Evolution of Rosaceae Fruit Types Based on Nuclear Phylogeny in the Context of Geological Times and Genome Duplication.” Molecular Biology and Evolution 34 (2): 262–81.

Xiong, Wenwei, Limei He, Jinsheng Lai, Hugo K. Dooner, and Chunguang Du. 2014. “HelitronScanner Uncovers a Large Overlooked Cache of Helitron Transposons in Many Plant Genomes.” Proceedings of the National Academy of Sciences of the United States of America 111 (28): 10263–68.

Xu, Zhao, and Hao Wang. 2007. “LTR_FINDER: An Efficient Tool for the Prediction of Full-Length LTR Retrotransposons.” Nucleic Acids Research 35 (Web Server issue): W265–68.

Yang, Z. 1997. “PAML: A Program Package for Phylogenetic Analysis by Maximum Likelihood.” Computer Applications in the Biosciences: CABIOS 13 (5): 555–56.

Yang, Z., and R. Nielsen. 2000. “Estimating Synonymous and Nonsynonymous Substitution Rates under Realistic Evolutionary Models.” Molecular Biology and Evolution 17 (1): 32–43.

Zhang, Huiting, Eric K. Wafula, Jon Eilers, Alex E. Harkess, Paula E. Ralph, Prakash Raj Timilsena, Claude W. dePamphilis, Jessica M. Waite, and Loren A. Honaas. 2022. “Building a Foundation for Gene Family Analysis in Rosaceae Genomes with a Novel Workflow: A Case Study in Pyrus Architecture Genes.” Frontiers in Plant Science 13 (November): 975942.

Zhang, Ming-Yue, Cheng Xue, Hongju Hu, Jiaming Li, Yongsong Xue, Runze Wang, Jing Fan, et al. 2021. “Genome-Wide Association Studies Provide Insights into the Genetic Determination of Fruit Traits of Pear.” Nature Communications 12 (1): 1144.

Zheng, Xiaoyan, Danying Cai, Daniel Potter, Joseph Postman, Jing Liu, and Yuanwen Teng. 2014. “Phylogeny and Evolutionary Histories of Pyrus L. Revealed by Phylogenetic Trees and Networks Based on Data from Multiple DNA Sequences.” Molecular Phylogenetics and Evolution 80 (November): 54–65.

Zhou, Chenxi, Shane A. McCarthy, and Richard Durbin. 2023. “YaHS: Yet Another Hi-C Scaffolding Tool.” Bioinformatics 39 (1). 10.1093/bioinformatics/btac808.

Chagné, David, Ross N. Crowhurst, Massimo Pindo, Amali Thrimawithana, Cecilia Deng, Hilary Ireland, Mark Fiers, et al. “The Draft Genome Sequence of European Pear (Pyrus Communis l. ‘Bartlett’).” PLoS ONE 9, no. 4 (April 3, 2014). 10.1371/journal.pone.0092644.

Linsmith, Gareth, Stephane Rombauts, Sara Montanari, Cecilia H Deng, Jean-Marc Celton, Philippe Guérif, Chang Liu, et al. “Pseudo-Chromosome–Length Genome Assembly of a Double Haploid ‘Bartlett’ Pear (Pyrus Communis L.).” GigaScience 8, no. 12 (December 9, 2019). 10.1093/gigascience/giz138.

Manni, M., Berkeley, M. R., Seppey, M., & Zdobnov, E. M. (2021). BUSCO: Assessing genomic data quality and beyond. Current Protocols, 1, e323. doi: 10.1002/cpz1.323

Manni, M., Matthew R Berkeley, Mathieu Seppey, Felipe A Simão, Evgeny M Zdobnov, BUSCO Update: Novel and Streamlined Workflows along with Broader and Deeper Phylogenetic Coverage for Scoring of Eukaryotic, Prokaryotic, and Viral Genomes. Molecular Biology and Evolution, Volume 38, Issue 10, October 2021, Pages 4647–4654

